# Context-dependent Requirements for FimH and Other Canonical Virulence Factors in Gut Colonization by Extraintestinal Pathogenic *Escherichia coli*

**DOI:** 10.1101/166108

**Authors:** Colin W. Russell, Brittany A. Fleming, Alan T. Stenquist, Morgan A. Wambaugh, Mary P. Bronner, Matthew A. Mulvey

**Author notes:** Address correspondence to: Matthew A Mulvey.

## Abstract

Extraintestinal pathogenic *Escherichia coli* (ExPEC) act as commensals within the mammalian gut, but can induce pathology upon dissemination to other host environments such as the urinary tract and bloodstream. It is thought that ExPEC genomes are shaped in large part by evolutionary forces encountered within the gut where the bacteria spend much of their time, provoking the question of how their extraintestinal virulence traits arose. The principle of coincidental evolution, in which a gene that evolved in one niche happens to be advantageous in another, has been used to argue that ExPEC virulence factors originated in response to selective pressures within the gut ecosystem. As a test of this hypothesis, the fitness of ExPEC mutants lacking canonical virulence factors was assessed within the intact murine gut in the absence of any antibiotic treatment. We found that most of the tested factors—including CNF1, Usp, colibactin, flagella, and the plasmid pUTI89—were dispensable for gut colonization. Deletion of genes encoding the adhesin PapG or the toxin HlyA caused transient defects, but did not affect longer-term persistence. In contrast, a mutant missing the type 1 pilus-associated adhesin FimH displayed reduced persistence within the gut. However, this phenotype was variable, being dependent on the presence of specific competing strains and partially attributable to aberrant flagellin expression by the *fimH* mutant. These data indicate that FimH and other key ExPEC-associated factors are not strictly required for gut colonization, suggesting that selective pressures within the gut do not drive the development of all extraintestinal virulence traits.

## INTRODUCTION

The bacterium *Escherichia coli* was first described by Theodor Escherich in 1884, and has since become a critical model organism that has been used to understand the fundamentals of molecular biology (1). *E. coli* is able to live in a variety of locations, including the soil, water, and the human gut. Although it is a prominent member of the neonatal microbiota, it is quickly overshadowed by burgeoning anaerobic bacteria as oxygen becomes scarce within the gut following birth (2, 3). In adults, *E. coli* is present at about 10^7^ to 10^9^ Colony Forming Units (CFU)/g feces, a level that is 100–10,000-fold-lower than the resident anaerobes (4). Despite being a minor component of the microbiota, the estimated number of *E. coli* cells that are transmitted via fecal matter from each human being to the environment in a single day is staggering – about 10^11^ CFU (5).

Understanding the role of *E. coli* within the microbiota is complicated by the fact that *E. coli* is a very diverse species with a wide spectrum of phenotypes (1). Some *E. coli* strains live harmlessly in the gut, while others act as pathogens, causing diarrhea and hemorrhaging (6). Others have been linked with the development of Crohn’s Disease and colorectal cancer (7–9). One strain, Nissle 1917, acts as a probiotic that assuages inflammation in addition to inhibiting colonization by pathogens such as *Salmonella* (10, 11). A group of strains known as Extraintestinal Pathogenic *Escherichia coli* (ExPEC) generally act as commensals within the gut, but can disseminate to other host environments and subsequently cause disease (12). ExPEC include uropathogenic *E. coli* (UPEC), which cause the overwhelming majority of urinary tract infections (UTIs) (13). These infections are especially prevalent among women, about half of whom will have at least one UTI during their lifetime. ExPEC is also responsible for other, more serious conditions, including sepsis and neonatal meningitis (12, 14).

The gut is thought to be the major ExPEC reservoir that seeds extraintestinal infections. Evidence for this notion is that the same ExPEC strain can often be isolated from both the feces and urine of individual patients suffering from UTIs (15–18). Indeed, ExPEC strains are frequently difficult to clear from the gut with antibiotic treatments, even when the pathogens are effectively eliminated from the urinary tract (19). Furthermore, *E. coli* strains belonging to phylogenetic group B2, which includes many ExPEC isolates, are much more likely to be long-term residents within the gut in comparison with other *E. coli* populations (20–22). The majority of adults carry group B2 *E. coli* strains within the gut, irrespective of extraintestinal infections (22). Cumulatively these observations suggest that ExPEC primarily inhabit the gut, with sporadic departures to extraintestinal sites.

Given that ExPEC reside mostly within the gut, and that transmission of ExPEC among individuals likely occurs chiefly through fecal-oral routes (4, 23–25), it is expected that ExPEC genomes have been shaped in large part by the evolutionary pressures present within the gut. How then, did extraintestinal virulence factors come into being? The hypothesis of coincidental evolution has often been evoked to answer this question (26). In general terms, coincidental evolution is when a factor evolves in one context, but happens to be useful in another context as well (27). When the hypothesis of coincidental evolution is applied to the ExPEC life cycle, the implication is that factors that promote virulence in extraintestinal niches evolved in the gut for a function possibly unrelated to virulence.

Little concrete evidence has been put forth to support or contradict coincidental evolution in the context of ExPEC infection, other than the fact that known extraintestinal virulence factors are often encoded by gut isolates (21, 26, 28). One prediction of this hypothesis is that extraintestinal virulence factors should play a role in gut colonization. To date, this possibility has only been addressed by one study in which an ExPEC mutant that lacks multiple pathogenicity-associated islands (PAIs) was found to be defective in its ability to persist within the murine intestinal tract (5). It is clear that more experimental work needs to be done to determine to what extent coincidental evolution applies to single virulence factors within ExPEC. To address this issue, several canonical virulence factors were individually deleted from a reference ExPEC isolate and the resulting mutants were tested for their ability to colonize and persist within the mouse gastrointestinal tract. Among eight virulence factors that were examined, only the type 1 pilus-associated mannose-binding adhesin FimH had a notable persistence defect within the gut. However, this defect was variable and partly contingent upon aberrant flagellin expression by the *fimH* mutant and the presence of specific competing strains. These findings are discussed in the context of both coincidental evolution and the development of anti-ExPEC therapeutics.

## RESULTS

### ExPEC stably colonizes the murine intestinal tract in the presence of the natural, intact microbiota

To examine ExPEC colonization of the intestinal tract, we employed a model in which adult specific pathogen free (SPF) Balb/c mice were intragastrically gavaged with ~10^9^ CFUs of the reference cystitis isolate F11. At various time points post-gavage, feces were collected, and the numbers of viable ExPE C present were enumerated. F11 and other bacterial strains used in this study were engineered to express either chloramphenicol (Clm^R^) or kanamycin (Kan^R^) resistance cassettes so that they could be easily identified by plating fecal homogenates on selective media. Following inoculation, the fecal titers of F11 remained fairly stable for up to 75 days, with median values ranging between 10^6^ and 10^7^ CFU/g feces after the first day (Fig. 1A). These data demonstrate that ExPEC can efficiently initiate and maintain colonization of the SPF mouse gut, in line with recent reports from our group and others (24, 29–31). Consistent with the observation that nonpathogenic *E. coli* mostly reside within the large bowel (32), the cecum and colon carried the largest load of F11 at the 2-week time point, although considerable numbers of F11 were also present within the small intestine (Fig. 1B).

**Figure 1.**
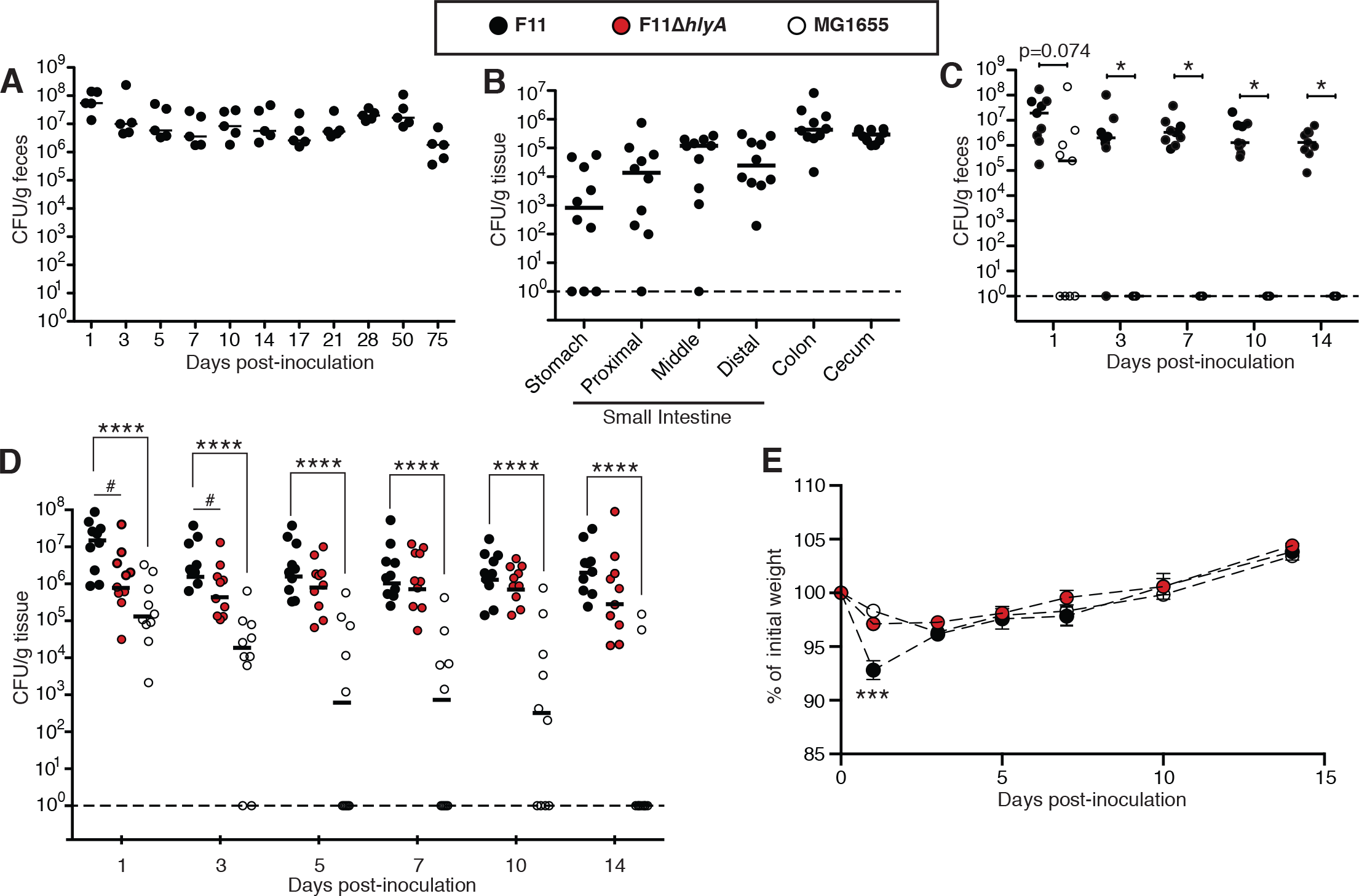
ExPEC colonizes and persists within the gut of SPF mice without causing serious pathology. Adult female SPF Balb/c mice were inoculated via oral gavage with ~10^9^ CFU of bacteria. (A) Titers of F11 recovered from the feces of mice at various time points post-gavage. n=5 mice. (B) F11 titers within intestinal tissues at 14 d post-gavage. (C) Mice were gavaged with a 1:1 mixture of F11 and MG1655, and fecal titers were determined for both populations at the indicated time points. *, *p* ≤ 0.05 by Wilcoxon signed-rank tests with corrections for multiple comparisons. (D) Fecal titers from non-competitive assays in which mice were orally inoculated with F11, F11Δ*hlyA*, or MG1655. ****, *p* ≤ 0.0001 by Mann-Whitney U Tests with corrections for multiple comparisons. #, *p* ≤ 0.05 by Mann-Whitney U Tests without corrections. In A-D, bars indicate median values. (E) Relative weights (mean values ± SD) of mice following oral inoculation with F11, F11Δ*hlyA*, or MG1655. Data were normalized to the mass of each mouse prior to gavage. ***, *p* ≤ 0.001 with corrections when F11 is compared to either F11Δ*hlyA* or MG1655 by unpaired Student’s *t* tests. In B-E, n=10 mice from two independent experiments. Adult female SPF Balb/c mice were inoculated via oral gavage with ~10^9^ CFU of bacteria. (A) Titers of F11 recovered from the feces of mice at various time points post-gavage. n=5 mice. (B) F11 titers within intestinal tissues at 14 d post-gavage. (C) Mice were gavaged with a 1:1 mixture of F11 and MG1655, and fecal titers were determined for both populations at the indicated time points. *, *p* ≤ 0.05 by Wilcoxon signed-rank tests with corrections for multiple comparisons. (D) Fecal titers from non-competitive assays in which mice were orally inoculated with F11, F11Δ*hlyA*, or MG1655. ****, *p* ≤ 0.0001 by Mann-Whitney U Tests with corrections for multiple comparisons. #, *p* ≤ 0.05 by Mann-Whitney U Tests without corrections. In A-D, bars indicate median values. (E) Relative weights (mean values ± SD) of mice following oral inoculation with F11, F11Δ*hlyA*, or MG1655. Data were normalized to the mass of each mouse prior to gavage. ***, *p* ≤ 0.001 with corrections when F11 is compared to either F11Δ*hlyA* or MG1655 by unpaired Student’s *t* tests. In B-E, n=10 mice from two independent experiments. Adult female SPF Balb/c mice were inoculated via oral gavage with ~10^9^ CFU of bacteria. (A) Titers of F11 recovered from the feces of mice at various time points post-gavage. n=5 mice. (B) F11 titers within intestinal tissues at 14 d post-gavage. (C) Mice were gavaged with a 1:1 mixture of F11 and MG1655, and fecal titers were determined for both populations at the indicated time points. *, *p* ≤ 0.05 by Wilcoxon signed-rank tests with corrections for multiple comparisons. (D) Fecal titers from non-competitive assays in which mice were orally inoculated with F11, F11Δ*hlyA*, or MG1655. ****, *p* ≤ 0.0001 by Mann-Whitney U Tests with corrections for multiple comparisons. #, *p* ≤ 0.05 by Mann-Whitney U Tests without corrections. In A-D, bars indicate median values. (E) Relative weights (mean values ± SD) of mice following oral inoculation with F11, F11Δ*hlyA*, or MG1655. Data were normalized to the mass of each mouse prior to gavage. ***, *p* ≤ 0.001 with corrections when F11 is compared to either F11Δ*hlyA* or MG1655 by unpaired Student’s *t* tests. In B-E, n=10 mice from two independent experiments. Adult female SPF Balb/c mice were inoculated via oral gavage with ~10^9^ CFU of bacteria. (A) Titers of F11 recovered from the feces of mice at various time points post-gavage. n=5 mice. (B) F11 titers within intestinal tissues at 14 d post-gavage. (C) Mice were gavaged with a 1:1 mixture of F11 and MG1655, and fecal titers were determined for both populations at the indicated time points. *, *p* ≤ 0.05 by Wilcoxon signed-rank tests with corrections for multiple comparisons. (D) Fecal titers from non-competitive assays in which mice were orally inoculated with F11, F11Δ*hlyA*, or MG1655. ****, *p* ≤ 0.0001 by Mann-Whitney U Tests with corrections for multiple comparisons. #, *p* ≤ 0.05 by Mann-Whitney U Tests without corrections. In A-D, bars indicate median values. (E) Relative weights (mean values ± SD) of mice following oral inoculation with F11, F11Δ*hlyA*, or MG1655. Data were normalized to the mass of each mouse prior to gavage. ***, *p* ≤ 0.001 with corrections when F11 is compared to either F11Δ*hlyA* or MG1655 by unpaired Student’s *t* tests. In B-E, n=10 mice from two independent experiments.

It is of note that the SPF mice utilized in these experiments were not treated with antibiotics and therefore each possesses an intact microbiota. This is in contrast to other commonly used mouse models in which mice are treated with streptomycin and/or other antibiotics in order to disrupt the intestinal microbiota and open up niches that can then be occupied by incoming microbes (32). Since antibiotic treatment was not required for consistent colonization of the gut by F11 (or by other ExPEC isolates (24, 29)), we wondered whether ExPEC is simply more adept at gut colonization than nonpathogenic *E. coli* strains. As a test of this idea, we first competed F11 head-to-head with MG1655, an often-used nonpathogenic *E. coli* K12 strain. Following oral gavage with equal numbers of F11 and MG1655, the K12 bacteria were cleared from all of the mice by day 3, while F11 stably persisted (Fig. 1C). When mice were gavaged with MG1655 alone, it was lost at a slower rate, but was still cleared from 80% of the mice by day 14 (Fig. 1D). This is in sharp contrast to the persistent phenotype observed with F11. Similar results were obtained using SPF C57Bl/6 mice (Fig. S1A).

To assess whether ExPEC colonization of the colon is marked by inflammation (colitis), the mice were weighed at several time points after colonization, as weight loss frequently accompanies colitis. Mice that were gavaged with F11 consistently experienced transient weight loss at the one-day time point, in comparison to mice that were inoculated with MG1655 (Fig. 1E). Since the ExPEC-associated pore forming toxin *α*-hemolysin (HlyA) was previously shown to induce inflammation within the gut in other mouse models (33–35), we hypothesized that HlyA contributes to the transient weight loss seen in F11-colonized animals. When mice were colonized with F11Δ*hlyA*, fecal titers of the mutant were notably lower at 1 and 3 days post-inoculation, relative to wild type (WT) F11 (Fig. 1D). However, the Δ*hlyA* mutant persisted much like the WT strain at later time points. These data, coupled with the observation that F11Δ*hlyA* does not elicit transient weight loss by the host (Fig. 1E), suggest that HlyA both enhances initial colonization of the gut by F11 and stimulates short-term inflammation that causes transient weight loss. By the 14-day time point, the colons of mice that were gavaged with WT F11, F11Δ*hlyA*, or MG1655 appear unperturbed, having normal crypt architecture and no evidence of infiltrating immune cells, as assessed by histological analysis. Altogether, these data indicate that F11 colonization induces transient inflammation of the intestinal tract with coordinate weight loss by the host. HlyA can facilitate these processes, but is not required for the longer-term persistence of F11 within the gut.

### Not all ExPEC-associated toxins promote ExPEC fitness within the gut

In light of the hypothesis of coincidental evolution and observations showing that HlyA can enhance initial colonization of the gut by F11 (Fig. 1D), we wished to examine possible roles for other canonical ExPEC-associated toxins as mediators of intestinal colonization. The first of these was cytotoxic necrotizing factor type 1 (CNF1), a secreted toxin that catalyzes the deamidation of a specific glutamine residue within Rho family GTPases (36). This causes constitutive activation of Rho GTPases, leading to aberrant host cytoskeletal rearrangements and multinucleation (2, 37, 38). CNF1 has been linked with ExPEC strains in epidemiological studies (38), and can enhance host inflammatory responses and ExPEC virulence in mouse models of UTI (39) and prostatitis (40). However, the effects of CNF1 on ExPEC fitness within the host remain unclear, clouded somewhat by conflicting reports (39–42). To test if CNF1 plays a role in the gut, mice were gavaged with 10^9^ CFU of either WT F11 or F11Δ*cnf1* and intestinal colonization levels were tracked by homogenizing and plating feces at various time points. Median bacterial levels for both the WT and mutant strains ranged between 10^5^ and 10^6^ CFU per gram of feces over the course of two weeks (Fig. 2A). At no point were the WT and F11Δ*cnf1* titers significantly different, suggesting that CNF1 does not impact the ability of F11 to colonize the mouse gut.

**Figure 2.**
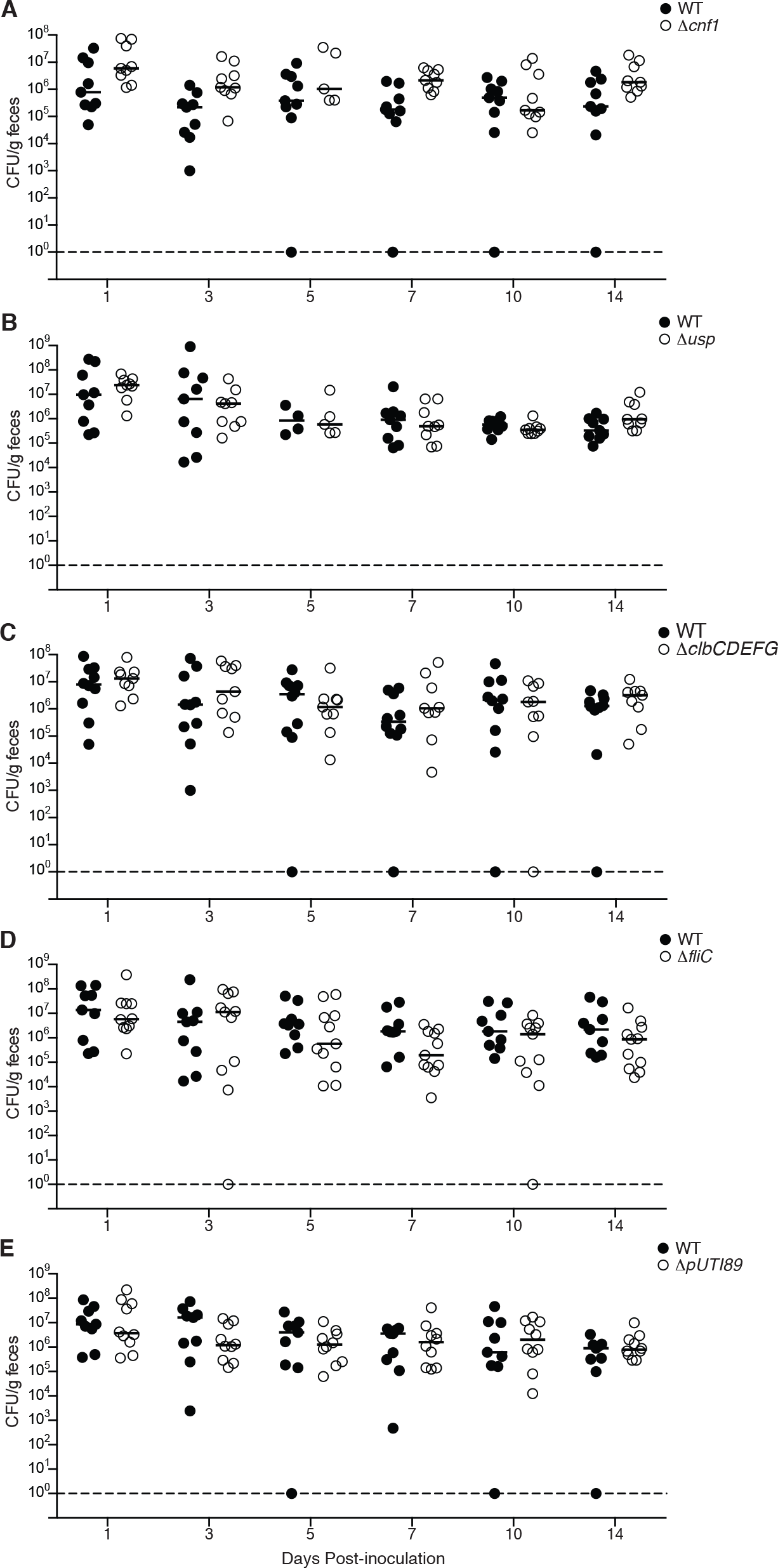
Key ExPEC-associated factors are not required for gut colonization. Mice were orally gavaged with ~10^9^ CFU of WT F11 or isogenic mutant strains lacking (A) *cnf1*, (B) *usp*, (C) *clbCDEFG*, (D) *fliC*, or (E) plasmid p*UTI89*. At the indicated time points, feces were collected, homogenized, and plated on selective media to determine bacterial titers. Bars represent median values. n=9–11 mice from two independent experiments. No statistically significant differences between WT and mutant titers were discerned using Mann-Whitney U tests.

Like HlyA and CNF1, Uropathogenic specific protein (Usp) is often encoded by ExPEC isolates (43, 44). Usp is also associated with *E. coli* fecal isolates that are capable of long-term persistence within the human infant intestinal tract (45). Interestingly, Usp has genotoxic nuclease activity as well as homology to colicins, a group of toxins that can be used by bacteria to harm competing microbes (46–48). Given that interbacterial competition within the gut is commonplace, we tested whether Usp is important for ExPEC gut colonization. The F11Δ*usp* mutant did not exhibit any notable defects within the gut, indicating that Usp is not required in this niche (Fig. 2B).

Another toxin that has been linked to ExPEC pathogenesis is colibactin, which is produced by a number of factors encoded by the polyketide synthase (*pks*) genomic island. The *pks* island is not typically carried by intestinal pathogenic *E. coli* strains, but is enriched in extraintestinal isolates relative to commensal fecal isolates (49). Colibactin induces DNA damage and cell cycle arrest in host cells (49, 50), and the presence of colibactin-producing bacteria has been linked with the development of colorectal cancer (51). In extraintestinal infections, colibactin-deficient bacterial strains are not as virulent as their WT counterparts. Mutation of the *pks* island reduces ExPEC translocation from the gut in neonatal rats (52) and reduces lymphopenia in septic mice (53). Whether the *pks* island also plays a role in bacterial fitness during gut colonization is not clear. Whereas one study observed decreased colonization of the small intestines of neonatal rats by colibactin mutants (52), another found no colonization differences between a *pks* mutant and WT bacteria (54). To test whether colibactin production is important in our adult mouse model of gut colonization, we orally inoculated mice with either WT F11 or F11Δ*clbCDEFG*, which lacks a key operon within the *pks* island (55). There was no difference in the colonization ability of the WT and mutant strains (Fig. 2C), suggesting that colibactin biosynthesis does not contribute to bacterial fitness in this model of gut colonization.

### Flagella are not required for ExPEC gut colonization

Flagella are filamentous organelles comprised of polymers of the flagellin protein FliC (56). Though best known for their role in motility, flagella can also promote bacterial attachment and biofilm formation, and can potently stimulate host inflammatory responses (57, 58). Several studies have provided evidence that flagella enhance ExPEC colonization of the mouse urinary tract (59–62). In contrast, flagella are not critical for gut colonization by nonpathogenic *E. coli* in streptomycin-treated mice (63). To determine if flagella are required for gut colonization by ExPEC in the face of an intact microbiota, we inoculated adult Balb/c mice with WT F11 or an isogenic mutant lacking *fliC*. In these assays, no clear differences between the WT strain and F11Δ*fliC* were observed (Fig. 2D), suggesting that flagella do not promote ExPEC fitness in the gut.

### The pUTI89 plasmid is not important for gut colonization

Many ExPEC isolates, including F11, carry plasmids that are identical or closely related to the pUTI89 plasmid that was first identified in the cystitis isolate UTI89 (64, 65). This plasmid, which is roughly 114 kb in length, encodes conjugation machinery, several virulence factors, and numerous hypothetical genes. Loss of pUTI89 impairs the fitness and intracellular growth of UTI89 during the early stages of a UTI in adult mice (66). Likewise, deletion of a closely related plasmid from the neonatal meningitis *E. coli* isolate RS218 attenuates bacterial virulence during systemic infections in rat pups (65). In addition, a number of pUTI89-associated genes have been linked with ExPEC mucus and glucose metabolism *in vitro* (30). To determine if pUTI89 facilitates gut colonization by F11, an F11 derivative that was cured of the plasmid was tested in our mouse model. The cured strain exhibited no defect when compared to the WT control strain (Fig. 2E), suggesting that pUTI89 is dispensable for ExPEC fitness within the gut.

### Disruption of *fimH* reduces gut colonization fitness, whereas the lack of *papG* has no effect

ExPEC strains encode many types of hair-like adhesive organelles known as pili, or fimbriae. Two of the most often-studied ExPEC-associated adhesins are P and type 1 pili (T1P) (67). P pili terminate with the PapG adhesin, which binds host globoseries glycosphingolipids and can facilitate bacterial infection of the kidneys. T1P are capped by the mannose-binding adhesin FimH, which promotes biofilm formation as well as bacterial attachment to and invasion of bladder epithelial cells. In our non-competitive gut colonization assays in which the WT and mutant bacterial strains are kept separate, the deletion of either *papG* or *fimH* did not significantly impair the ability of F11 to colonize the intestinal tract (Fig. 3A and B). However, in these assays we noted that the Δ*fimH* mutant was cleared in 3 out of 10 mice–more than observed with any of the other tested mutants, though not statistically significant (Fig. 3B). These results prompted us to examine the *fimH* mutant further using competitive assays in which the WT and mutant strains are inoculated as a 1:1 mixture into mice via oral gavage. In these and other competitive assays, total ExPEC levels remain fairly steady over time at around 10^6^ CFU/g feces, even when ratios of the individual competing ExPEC strains are variable.

**Figure 3.**
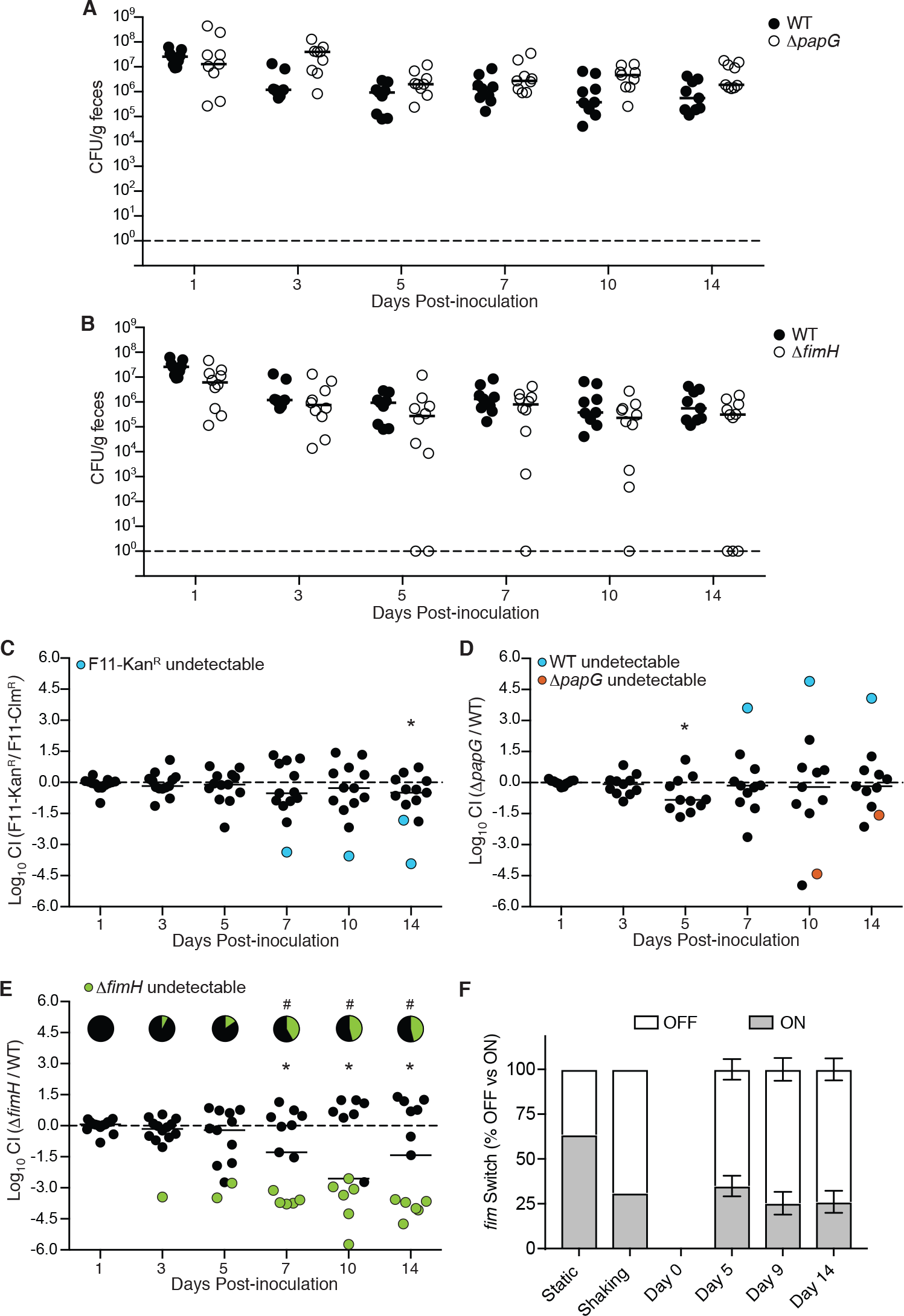
The persistence of F11 Δ*fimH*, but not F11 Δ*papG*, within the gut is impaired in competitive assays. Mice were gavaged with WT F11, (A) F11Δ*papG*, or (B) F11Δ*fimH* and fecal titers were determined at the indicated time points. In these non-competitive assays, no significant differences were identified by Mann-Whitney U tests. For competitive assays, mice were gavaged with a 1:1 mixture of (C) F11-Kan^R^ and F11-Clm^R^, (D) WT F11 and F11Δ*papG*, (E) or WT F11 and F11Δ*fimH*. *, *p* ≤ 0.05 by one sample *t* tests without correction for multiple comparisons. Pie charts in (E) indicate fraction of mice in which F11Δ*fimH* (green) was not detected. #, *p* ≤ 0.05 by Fisher’s exact tests without corrections. In A-E, n = 11–13 mice from two independent assays. (F) Graph shows fractions of the *fim* switch in the ON and OFF positions from fecal samples recovered from mice following oral inoculation with F11. Bars indicate mean values ± SEM; n=5–10 mice. Results from F11 grown in static or shaking LB broth are shown for comparison.

As a control, we first competed Kan^R^-and Clm^R^-tagged F11 strains against one another to assess if the resistance cassettes alone compromised bacterial fitness within the intestinal tract. In these control assays, F11-Kan^R^ exhibited a modest defect, amounting to about a 3-fold decrease relative to F11-Clm^R^ on day 14, with a median competitive index (CI; see Methods) of −0.49 (Fig. 3C). Likewise, no striking differences between F11-Clm^R^ and F11Δ*papG* (Kan^R^) were observed in competitive assays, with the exception of a transient 6.8-fold decrease (median CI of −0.83) in prevalence of the *papG* mutant on day 5 (Fig. 3D). In contrast, when F11Δ*fimH* (Kan^R^) was competed against F11-Clm^R^, the Δ*fimH* mutant became progressively worse than the control strain starting at day 7 post-inoculation (Fig. 3E). By day 10, there was about a 360-fold reduction in the relative levels of F11Δ*fimH* recovered in the feces, reflecting a median CI of −2.55. In addition, the Δ*fimH* mutant began to be cleared as early as day 3, and was not detected in the feces of nearly half of the mice by day 10 (green, Fig. 3E). In comparison, F11Δ*papG* dropped below levels of detection in only one mouse during the 14-d time course of these competitive assays (red, Fig. 3D). Of note, the differences between strains indicated in Figs. 3C-E are not statistically significant when *p* values are corrected for multiple comparisons. However, the amplitude (effect size) of the phenotypes observed with the Δ*fimH* mutant suggests that disruption of this gene can markedly impair ExPEC persistence within the gut in competitive assays.

### T1P expression by F11 is modest following excretion from the gut

T1P are phase variable, being turned ON or OFF through recombinase-mediated flipping of an invertible promoter element within the *fim* gene cluster (67). The orientation of this *fim* switch can be monitored and quantified by PCR as a means to assess levels of T1P expression (68, 69). Within the feces of mice that are colonized by F11, we found that the *fim* switch is in the ON position in about 25 to 35% of the excreted bacteria recovered on days 5, 9, and 14 post-inoculation (Fig. 3F). This is on par with results from shaking broth cultures, in which the *fim* switch is skewed towards the OFF position. These data indicate that T1P expression by feces-associated ExPEC is limited, which may enable the pathogen to better disseminate either within the intestinal tract or after being discharged from the host.

### Increased flagellin expression partially explains the colonization defect observed with F11Δ*fimH*Δ

T1P, and FimH in particular, may enhance ExPEC persistence within the gut via multiple mechanisms, such as aiding the formation of protective biofilm-like communities or facilitating bacterial attachment to intestinal tissues (58, 70, 71). However, it is conceivable that disruption of *fimH* could also reduce ExPEC fitness within the gut via effects on other bacterial or host processes. Specifically, previous work showed that deletion of the entire *fim* operon in the ExPEC isolate UTI89 causes the upregulation of FliC with a coordinate increase in swimming motility (72). This could be problematic for *fim* mutants since it is known that aberrant overexpression of FliC can impair ExPEC colonization of the gut (5), possibly as a consequence of FliC-mediated stimulation of host inflammatory responses (57). Furthermore, within the intestinal tracts of germ-free or streptomycin-treated mice, mutations that reduce bacterial motility and FliC expression are selected and can promote the persistence of the K12 strain MG1655 (73–75). Together, these observations led us to question if defects in the ability of F11Δ*fimH* to survive within the gut in competitive assays are associated with altered FliC expression.

In assessing this possibility, we first noted that deletion of *fimH* does increase the motility of F11 in swim agar plates (Fig. 4A). This phenotype correlates with augmented FliC expression (Fig. 4B), as measured by use of the low-copy reporter construct p*fliC-lux* in which the *luxCDABE* gene cluster encoding bacterial luciferase is transcriptionally fused with the conserved *fliC* promoter (62). These results mirror those reported for a UTI89 mutant lacking the entire *fim* operon (72). Interestingly, in our assays the deletion of *papG* in F11 had nearly the opposite effect of the *fimH* deletion, greatly reducing the motility of F11 and ablating FliC expression (Fig. 4A-B).

**Figure 4.**
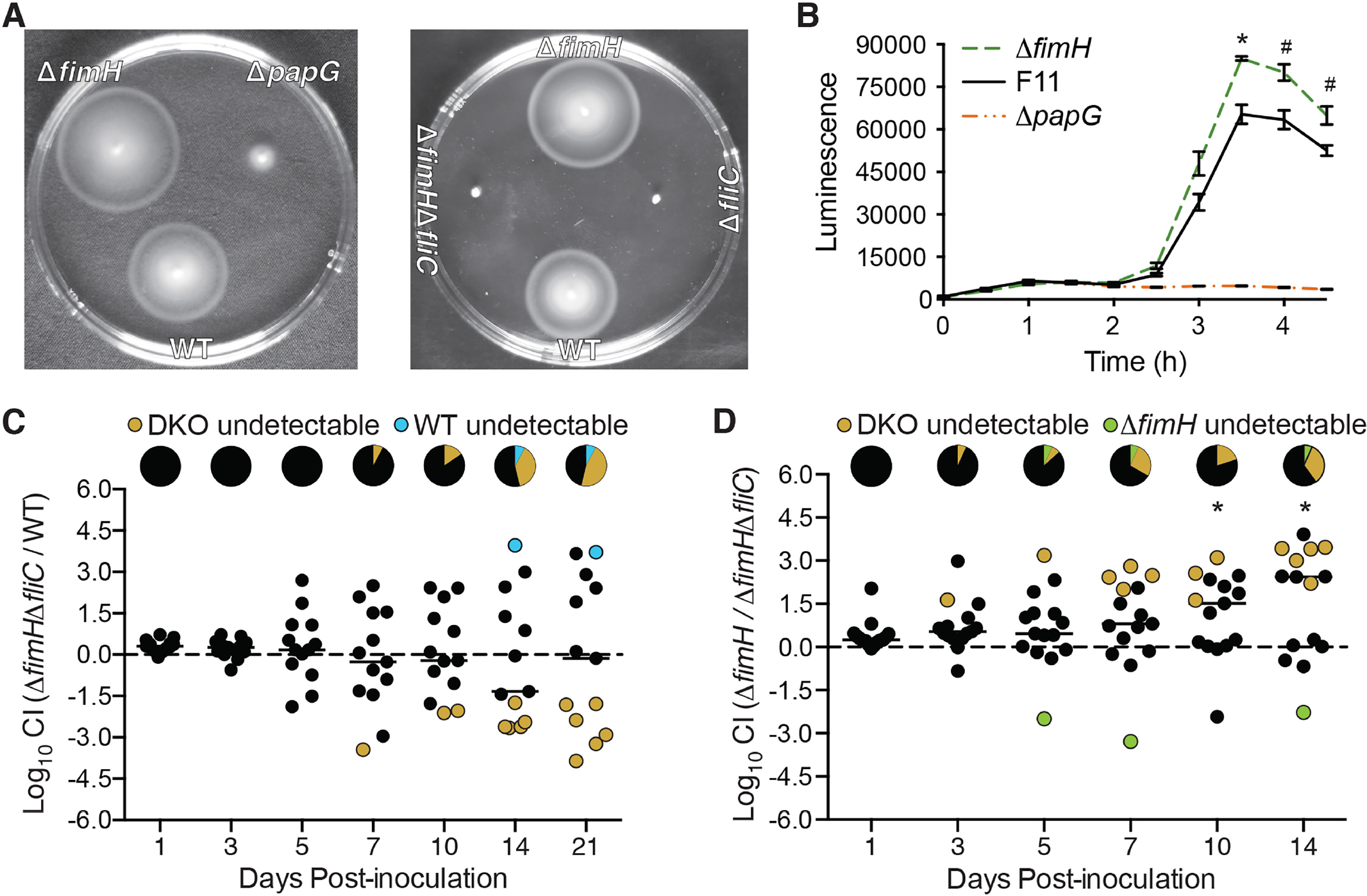
Flagellin expression impacts the efficacy of gut colonization by F11Δ*fimH*. (A) WT F11 and the indicated mutant derivatives were inoculated into motility agar to assess swimming. Images of swim plates were taken at 8–10 h post-inoculation and are representative of results from three independent assays. (B) Plot shows results from *fliC* expression reporter assays with WT F11, F11Δ*fimH*, and F11Δ*papG* carrying p*fliC-lux*. Lines indicate mean luminescence values ± SEM from three independent assays performed in triplicate. *p* ≤ 0.05 by multiple *t* tests with (*) or without (#) corrections for multiple comparisons. (C-D) Graphs show results from competitive assays in which mice were orally gavaged with a 1:1 mixture of (C) WT F11 and F11Δ*fimH*Δ*fliC* (DKO) bacteria or (D) F11Δ*fimH* and F11Δ*fimH*Δ*fliC* (DKO). Fecal titers of each strain were determined at the indicated time points by plating on selective media. *, *p* ≤ 0.05 by one sample *t* tests with corrections for multiple comparisons. Pie charts in (C-D) denote the fraction of mice in which the DKO mutant (yellow), WT F11 (blue), or F11Δ*fimH* (green) were not detected. No significant differences were discerned by Fisher’s exact tests. n=13–15 mice from at least two independent assays.

To test if FliC contributes to the defects in gut colonization observed with F11Δ*fimH*, we generated a double knockout (DKO) mutant lacking both *fimH* and *fliC*. This mutant (F11Δ*fimH*Δ*fliC*) is non-motile in swim plates, similar to the single Δ*fliC* mutant strain (Fig. 4A). In competitive gut colonization assays with WT F11, the Δ*fimH*Δ*fliC* mutant exhibited less pronounced deficiencies than F11Δ*fimH* (Fig. 4C; compare with Fig. 3E). The greatest defect was observed at day 14 post-inoculation, at which point F11Δ*fimH*Δ*fliC* was not detected in the feces of 5 of the 13 mice. Relative to the WT strain, F11Δ*fimH*Δ*fliC* titers were reduced by about 21-fold on day 14, corresponding with a median CI of −1.33. This defect was substantially smaller than the maximal ~360-fold reduction seen with the single Δ*fimH* mutant in competition with WT F11 on day 10 post-inoculation (see Fig. 3E). In addition, clearance of F11Δ*fimH*Δ*fliC* was not observed until day 7 post-inoculation (Fig 4C), whereas loss of F11Δ*fimH* was evident starting at day 3 (Fig 3C). In light of the delayed defects seen with F11Δ*fimH*Δ*fliC*, we extended the assays to day 21. At this point, the median CI value returned close to 0, though F11Δ*fimH*Δ*fliC* was still undetectable in feces from nearly half of the mice while the WT strain was present in all but one sample (Fig. 4C). Differences between F11Δ*fimH*Δ*fliC* and the WT strain were not statistically significant, either with or without corrections for multiple comparisons. The less conspicuous defects seen with F11Δ*fimH*Δ*fliC* in these assays suggest that aberrant FliC expression does contribute to the compromised persistence of F11Δ*fimH* within the gut.

### F11Δ*fimH* outcompetes F11Δ*fimH*Δ*fliC* within the gutΔ

To better understand the effects that the loss of *fimH* and *fliC* have on bacterial fitness during gut colonization, the Δ*fimH*Δ*fliC* and Δ*fimH* mutants were directly competed. We hypothesized that the Δ*fimH* strain, with heightened FliC expression, would be outcompeted by the Δ*fimH*Δ*fliC* DKO mutant. However, F11Δ*fimH*Δ*fliC* exhibited clear and statistically significant defects in comparison with F11Δ*fimH* (Fig 4D). Fecal titers of the DKO mutant were below levels of detection in one out of 15 mice on day 3 post-inoculation, and by day 14 the DKO mutant was undetectable in the feces of a third of the animals. At this point, F11Δ*fimH* was about 270-fold more abundant than the DKO mutant, corresponding with a median CI of 2.43. These data, as well results from non-competitive assays (see Fig. 3B), indicate that the importance of FimH to ExPEC survival within the intestinal tract is dependent upon the nature of the competing microbes that are present.

### WT F11 and F11Δ*fimH* can outcompete one other in reciprocal serial colonization assays

To further evaluate FimH requirements within the intestinal tract, we carried out serial colonization assays in which Balb/c mice were first orally gavaged with WT F11 before the introduction of F11Δ*fimH* 7 d later. Though fecal titers of the Δ*fimH* mutant were initially high and on par with the WT strain, levels of the mutant dropped precipitously and were below the limits of detection in most mice by day 3 post-inoculation (Fig. 5A). In reciprocal experiments, in which F11Δ*fimH* was allowed to colonize the mice prior to instillation of the WT strain, F11Δ*fimH* persisted while WT F11 was cleared from most of the mice by day 10 (Fig. 5B). These data show that FimH is not strictly required for ExPEC to prevent colonization of the gut by a new competing strain, though the adhesin does seem to aid this process. In addition, these results indicate that WT F11 and F11Δ*fimH* likely vie for the same intestinal niches, with the first strain established having the upper hand irrespective of FimH expression.

**Figure 5.**
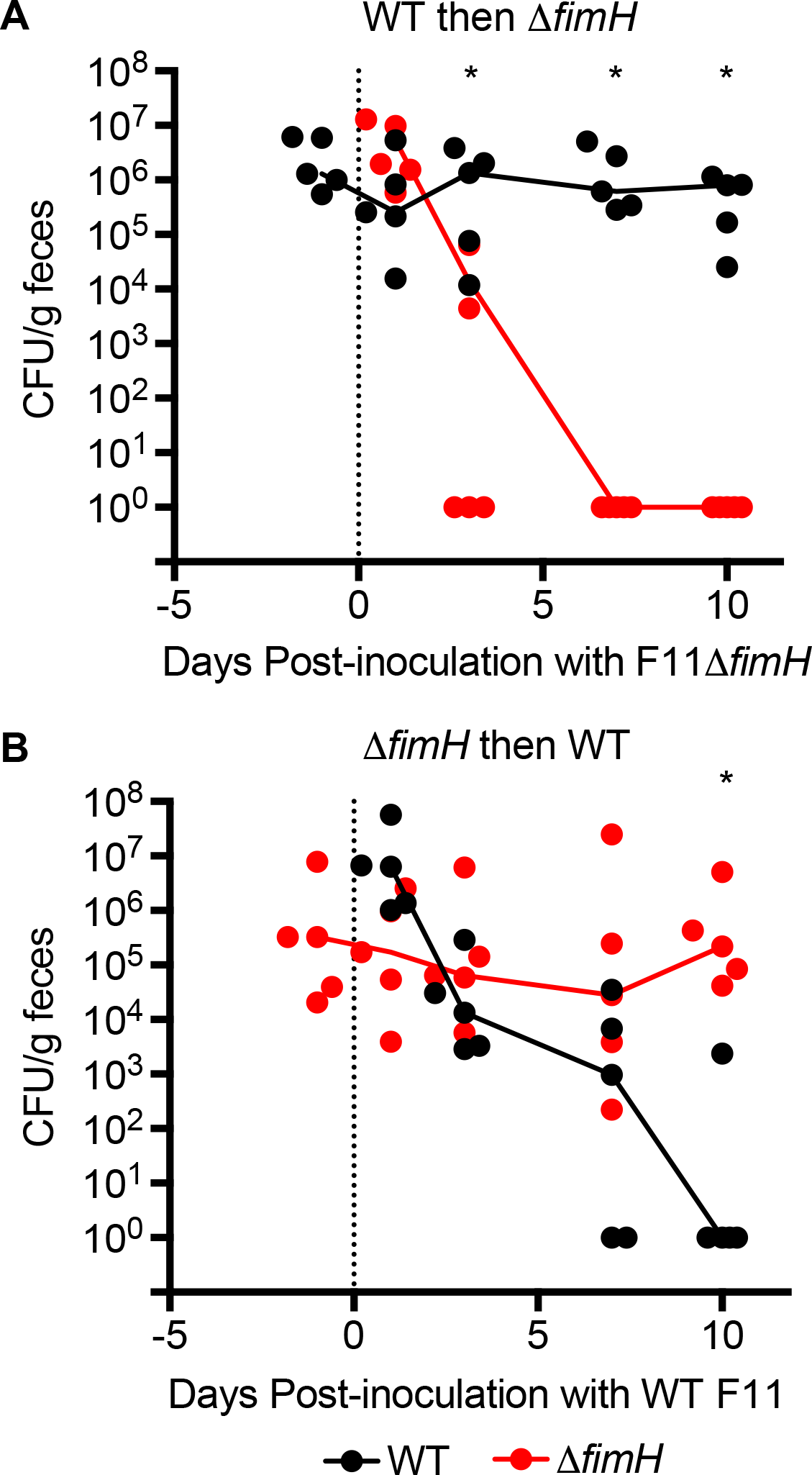
Pre-colonization of mice with F11*ΔfimH* effectively limits colonization by the WT strain, and vice versa. Balb/c mice were inoculated via oral gavage with (A) WT F11 (specifically F11-Clm^R^) and then with F11 *ΔfimH* (Kan^R^) 7 d later. (B) Alternatively, mice were inoculated with F11 *ΔfimH* followed 7 d later by the WT strain. Solid lines connect median fecal titers of each strain over time. The zero time point (dotted line) indicates when the second strain (WT F11 or F11*ΔfimH*) was introduced. *, *p*<0.05 by Mann-Whitney U tests with corrections for multiple measurements; n=5 mice.

## DISCUSSION

The concept of coincidental evolution, coupled with phylogenetic analyses, suggests that the extraintestinal success of ExPEC strains is a by-product of their ability to colonize the gut (26). A corollary of this hypothesis is that extraintestinal virulence and fitness factors promote ExPEC colonization of the gut. In support of this possibility, researchers previously showed that deletion of the seven major pathogenicity islands (PAIs) of the ExPEC isolate 536 not only reduced the virulence of this pathogen in a murine sepsis model, but also attenuated pathogen persistence within the gut (5). Here, we set out to determine if the principal of coincidental evolution could be applied to individual virulence and fitness determinants encoded by the reference ExPEC strain F11. In non-competitive assays using adult SPF mice, we observed no defects in intestinal colonization or the persistence of ExPEC mutants lacking CNF1, Usp, colibactin, flagellin, or the plasmid pUTI89 (see Fig. 2). Mutants that are missing either *papG* or *hlyA* exhibited transient colonization defects, but these did not affect longer-term ExPEC survival within the gut (Figs. 1D, 3A, and 3D). These results indicate that at least some ExPEC-linked genes can influence bacterial fitness within the gut, though the effects may be modest. Discerning more unequivocal phenotypes for individual ExPEC-associated loci within the gut can be complicated and context-dependent, as exemplified by the analysis of Δ*fimH* mutants.

In non-competitive assays, we observed no significant differences between WT F11 and F11Δ*fimH*, although the mutant was cleared from a few mice over the course of the 14-d experiments (Fig. 3B). This prompted us to test F11Δ*fimH* further using competitive assays with the WT strain. In these assays, differences between the Δ*fimH* mutant and WT strains were more distinct, but not significantly so if the data are corrected for multiple comparisons. Still, the fact that F11Δ*fimH* titers within the feces from nearly half the mice fell below detectable levels by day 10 post-inoculation suggests that FimH can promote ExPEC persistence within the gut (Fig. 3E). These findings are in line with recently published work showing that intestinal persistence of the ExPEC isolate UTI89 within streptomycin-treated mice is facilitated by FimH expression (76). This study, as well as other work with enteric *E. coli* pathogens (71, 77, 78), suggests that FimH can promote ExPEC interactions with intestinal epithelial cells. In our mouse models, we observed that F11 is localized primarily within the lumen of the colon, though substantial numbers of the pathogen are also associated with the colonic tissue (see Fig. S1B). While FimH may mediate interactions between ExPEC and the intestinal epithelium (76), the adhesin could also influence pathogen persistence within the gut via effects on other processes, including biofilm development and the modulation of innate host defenses (58, 70, 79, 80).

Defining the contributions of FimH to ExPEC fitness within the gut is further complicated by the observation that the deletion of *fimH* enhances FliC expression by F11 and increases motility (Fig. 4). Analysis of a *fliC* and *fimH* DKO mutant in competition with WT F11 indicated that aberrant FliC expression is at least partially responsible for the colonization defects observed with the Δ*fimH* mutant (Figs. 3E and 4C). This situation mirrors results reported for the ExPEC strain 536, in which reduced intestinal persistence of the mutant lacking seven PAIs was attributed to FliC overexpression (see above, (5)). The expression of flagella is generally thought to have mostly detrimental effects on *E. coli* fitness within the gut, possibly due to an increased burden on bacterial metabolism and the ability of FliC to stimulate host inflammatory pathways (57, 73–75). However, in competitive assays F11Δ*fimH* outperformed F11Δ*fimH*Δ*fliC*, even though the single mutant is hypermotile (Fig. 4D). One potential explanation for this finding is that flagella expression might at times be an advantage for *E. coli* within one or more intestinal niche, as previously suggested (75). This possibility is difficult to reconcile with the observation that F11Δ*fliC* has no substantial defects in our noncompetitive assays (Fig. 2D). It is plausible that crosstalk among bacterial regulators of motility, T1P expression, and other adhesins also contribute to the phenotypes observed in our assays with the *fimH* mutants and other F11 derivatives (81–85), but this will require additional studies to tease apart.

The ability of distinct types of bacteria to utilize different spatial and nutritional niches within the gut allows for the coexistence of the diverse organisms that comprise the intestinal microbiota, and helps provide a barrier against colonization by newly arriving microbes (86). This latter effect, known as colonization resistance, is one reason that it is generally necessary to treat mice with an antibiotic like streptomycin to open up niches for incoming bacteria that are delivered into the gastrointestinal tract by oral gavage (32). A striking feature of our experimental system is that F11 is able to colonize and persist indefinitely within the intestinal tract of conventional SPF mice without the need to first administer antibiotics (Fig. 1A). Our group and others have recently reported similar results with distinct ExPEC isolates in different mouse strain backgrounds (24, 29, 31, 87). ExPEC may be able to effectively colonize our untreated mice because they have very low numbers of endogenous *E. coli* and other Enterobacteriaceae based on 16S rRNA gene sequencing and selective plating assays (CR Russell and MA Mulvey, unpublished data). Nevertheless, these animals are still resistant to colonization by the K12 strain MG1655, indicating that MG1655 lacks one or more genes that ExPEC employs to persist within the gut. When in competition with F11, MG1655 is cleared from the gut at a notably faster rate than in noncompetitive assays (see Fig. 1), suggesting that these two strains compete for the same intestinal niches.

A better understanding of the survival mechanisms used by ExPEC within the intestinal tract may aid the development of more efficacious probiotics, while also elucidating new therapeutic strategies to combat ExPEC before it is able to disseminate and cause disease at sites beyond the intestinal tract. The effectiveness of such approaches may be dependent on multiple variables, including timing, the makeup of the microbiota, and the presence of specific competing strains that can alter ExPEC requirements for individual fitness determinants. For example, in contrast to the situation in competitive assays, F11Δ*fimH* has no trouble colonizing the gut in noncompetitive experiments and, once established, this mutant can even persist when challenged with the WT strain (see Figs. 3B and 5B). Thus, while FimH can facilitate ExPEC persistence within the gut in some settings, it is not always an absolute requirement. This conclusion may help reconcile results from older studies in which the expression of T1P was found to be unnecessary for *E. coli* persistence within the intestinal tracts of rodents and human infants (88–90).

Context-dependent variability in the phenotypic effects of fitness determinants like FimH may complicate treatment approaches, as well as our ability to discern how life within the gut affects the evolution of ExPEC virulence traits. Nevertheless, data presented here indicate that it is reasonable to apply the principle of coincidental evolution to some ExPEC-associated genes, though the phenotypic effects of these genes may be modest and variable dependent on the experimental system that is used. The lack of easily discernable phenotypes for ExPEC-associated loci within the gut suggests that selective forces encountered within other niches have also helped shape ExPEC genomes. For example, extraintestinal virulence has been correlated with the ability of ExPEC strains to resist killing by amoebae (91), while T1P expression has been linked with the transmission of ExPEC between individuals by promoting transient colonization of the oropharynx (88). In total, the results presented here show that piecing together the evolutionary history of ExPEC virulence and fitness traits is a complicated task. However, continuing efforts to resolve this problem will provide a more detailed picture of ExPEC ecology and may help identify niche-specific fitness determinants that could be attractive targets for therapeutic intervention.

## MATERIALS AND METHODS

### Bacterial strains

The cystitis isolate F11 and the K-12 strain MG1655 were genetically modified using lambda-red recombination that was facilitated by the pKM208 plasmid (92). Most of the constructs used for recombination were created by PCR using either pKD4 or pKD3 as a template to amplify a kanamycin or chloramphenicol resistance cassette, respectively, flanked by ~40 base pairs of DNA with homology to the target insertion site. In some cases, longer homology regions were required, and three-part PCR was performed. This was done by PCR amplification of an antibiotic resistance cassette and regions that are upstream and downstream of the target gene. Primers used were designed to contain sections of homology to allow the three PCR products to be stitched together in a single ligation reaction.

The pCP20 plasmid was used to remove the resistance cassette as necessary (93). To cure F11 of the pUTI89 plasmid, the *ccdAB* toxin-antitoxin system was replaced with a *tetA-sacB* construct and spontaneous loss of the plasmid was selected for on LB agar plates containing fusaric acid and sucrose, as explained previously (94). The strains used in this study are listed in Table S1 along with the primers used to create them. Prior to lambda-red recombination, bacteria were grown shaking in LB broth at 37ºC. All growth in petri dishes was done using LB agar supplemented with chloramphenicol (20 µg/ml), kanamycin (50 µg/ml), or ampicillin (50 µg/ml), as appropriate.

### Mouse gut colonization

Mice were handled in accordance with protocols approved by the Institutional Animal Care and Use Committee at the University of Utah (Protocol number 15-12015), following US federal guidelines indicated by the Office of Laboratory Animal Welfare (OLAW) and described in the Guide for the Care and Use of Laboratory Animals, 8th Edition.

Prior to inoculation into mice, bacteria were grown statically from frozen stocks for 24 h at 37ºC in 250 ml flasks containing 20 ml of modified M9 media [MgSO_4_•7H_2_O (1 mM), CaCl_2_•2H_2_O (0.1 mM), D-(+)-glucose (0.1%), nicotinic acid (0.00125%), thiamine HCl (0.00165%), casamino acids (0.2%), Na_2_HPO_4_ (6g/L), KH_2_PO_4_ (3 g/L), NH_4_Cl (1 g/L), and NaCl (0.5 g/L) in water]. A total of 12 ml of culture (6 ml of each culture for competitive experiments) was centrifuged at 8,000 X g for 10 min. Bacterial pellets were then washed once and resuspended in 0.5 ml of PBS. To inoculate the mouse gastrointestinal tract, 7–8 week old female, specific pathogen free (SPF) Balb/c or C57Bl/6 mice (The Jackson Laboratory) were orally gavaged with 50 µl PBS containing ~10^9^ CFU of bacteria. At various time points post-inoculation, individual mice were briefly placed into unused takeout boxes for weighing and feces collection. Feces were homogenized in 1 ml of 0.7% NaCl and then samples were briefly centrifuged to pellet insoluble debris. Supernatants were serially diluted and spread onto LB agar plates containing either chloramphenicol (20 µg/ml) or kanamycin (50 µg/ml) to select for growth of the relevant bacterial strains. Prior to gavage, fecal samples were analyzed to ensure that there were no endogenous bacteria present that were resistant to chloramphenicol or kanamycin. Mice were housed 3–5 per cage, and were allowed to eat (irradiated Teklad Global Soy Protein-Free Extruded chow) and drink antibiotic-free water *ad libitum*. Competitive indices (CIs) were calculated as the ratio of mutant over WT bacteria recovered in the feces divided by the ratio of mutant over WT bacteria in the inoculum, as noted previously (30).

To determine F11 (specifically F11-Clm^R^) titers within the various regions of the gastrointestinal tract at 14 d post-inoculation, mice were anaesthetized via isofluorane, euthanized by cervical dislocation, and the cecum, colon, small intestine, and stomach were removed. The small intestine was divided into thirds, with the portion closest to the stomach labeled “proximal” and the portion closest to the cecum labeled “distal”. A part of each organ was weighed and placed into a Safe-Lock tube (Eppendorf) with three 3.2 mm stainless steel beads, and homogenized using the Bullet Blender (Next Advance). The homogenates were then serially diluted and plated onto LB agar plates containing chloramphenicol (20 µg/mL) to quantify the levels of F11-*clmR* present.

For histology, colon tissues were fixed in 10% neutral buffered formalin and submitted to the University of Utah Research Histology core for processing and staining with hematoxylin and eosin (H&E). Uninfected intestinal tissues and tissues recovered from mice at 14 d post-inoculation with either MG1655 or F11Δ*hlyA* were used for comparison. Random colon tissue sections from 5 or more mice in each experimental group were analyzed in a blinded fashion by a trained pathologist (M.P. Bronner).

### Motility Assays

To test the swimming ability of particular strains, motility agar plates were made by pouring 25 mL of LB soft agar (0.1% agar) into a petri dish. Bacteria (1.5 µl from an overnight culture) were dispensed just below the surface of the plate, which was then incubated at 37º C for 8–10 h prior to imaging with a Stratagene Eagle Eye II System.

### Luciferase assays

The p*fliC-lux* construct was kindly provided by the Mobley lab (62). Overnight shaking cultures of F11, F11Δ*papG*, and F11Δ*fimH* carrying p*fliC-lux* were diluted 1:100 into 1 ml of fresh tryptone broth (Fisher Scientific) containing ampicillin (50 µg/ml). Three 100-ul-aliquots of each culture were then transferred to a white 96-well polystyrene plate (Dynex Technologies) and grown statically at 37°C. Luminescent emission spectra were collected every 30 minutes for 4.5 hours using Gen5 Software with a BioTek Synergy H1 plate reader. The instrument was set to integrate readings over 10 seconds using a gain of 135 and height of 1 mm. Before each reading, the plates were shaken for 1 second. Corresponding growth curves, which were obtained by taking OD_600_ measurements of cultures grown statically in clear 96-well polystyrene plates (CytoOne), showed that the strains used in these assays grew similarly.

### Analysis of the fim switch

Quantification of the *fim* switch in the ON and OFF positions was DNA was carried out essentially as previously described (68, 69). DNA was purified from feces using the ZR Fecal DNA MiniPrep kit, whereas DNA was purified from broth cultures using the Promega Wizard Genomic DNA Purification kit. To skew the *fim* switch towards the ON position, F11 was grown statically at 37°C in 20 mL LB broth in 250 mL Erlenmeyer flasks for 24 h, sub-cultured 1:1000 into fresh LB broth, and then incubated for another 24 h. To drive the *fim* switch towards the OFF position, bacteria were grown shaking in LB broth to exponential phase. The primers F11_fimS_F (TACCGCCAGTAATGCTGCTC) and F11_fimS_R (GTCCCACCATTAACCGTCGT) were used to amplify the *fim* switch region by PCR using the following thermocycler conditions: 95**°**C for 5 minutes, 10 cycles (95**°**C for 30 sec, 60**°**C for 20 sec, 72**°**C for 40 sec), followed by 30 cycles (95**°**C for 30 sec, 56**°**C for 20 sec, 72**°**C for 40 sec), and ending with a 5 min incubation at 72**°**C. The reaction products were column purified, digested with HinfI for 1 h at 37**°**C, resolved in 1% agarose gels, and imaged. Bands representing the *fim* switch in the ON and OFF positions were quantified using ImageJ.

### Statistical Analysis

All statistical tests were carried out using GraphPad Prism or Stata/IC-14 software. Where indicated, corrections for multiple comparisons were made using the Hochberg procedure (95). *P* values ≤ 0.05 were considered significant.

## FUNDING INFORMATION

This work was supported by NIH grant AI095647 and by seed grants provided by the University of Utah and the Pathology Department. The funders had no role in study design, data collection and interpretation, or the decision to submit the work for publication.

## ACKNOWLEDGMENTS

We are grateful to members of the Mulvey Lab for their insightful conversations and advice. We are also indebted to the Office of Comparative Medicine at the University of Utah for their assistance with animal care.

---

**Supplemental Figure S1.**
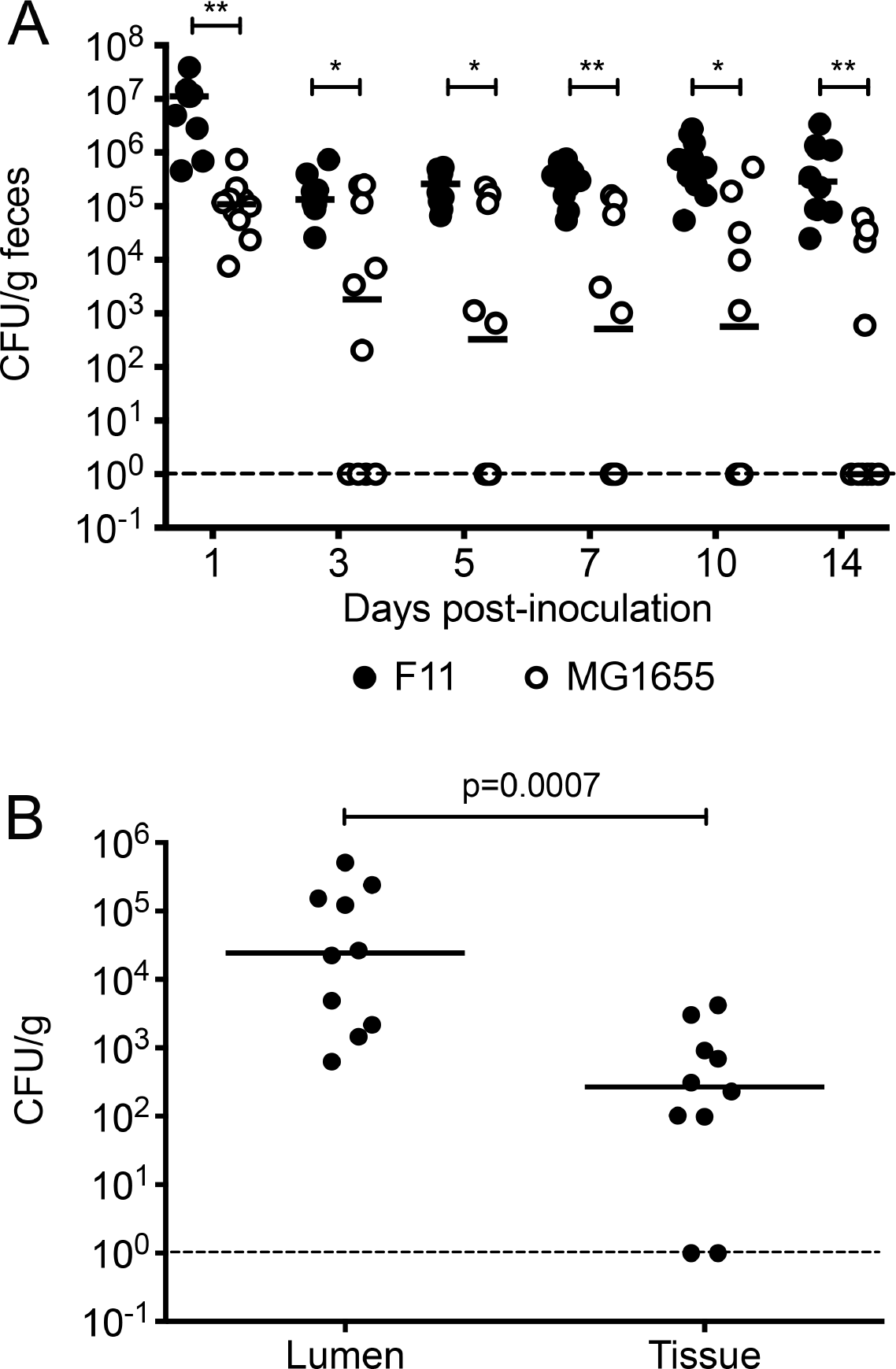
F11, but not MG1655, effectively colonizes the intestinal tract of SPF C57Bl/6 mice. Adult female SPF C57Bl/6 mice were inoculated via oral gavage with ~10^9^ CFU of either F11 or MG1655. (A) Graph shows titers of F11 and MG1655 recovered from the feces at various time points post-gavage. n=10 mice from two independent non-competitive assays. **, *p*<0.01 by Wilcoxon signed-rank tests, with corrections for multiple comparisons. (B) F11 titers recovered in association with the colonic tissue or within the lumen of the colon at 14 d post-gavage. *P* value determined by Mann-Whitney test; n=10 mice.

**Supplemental Table S1.**
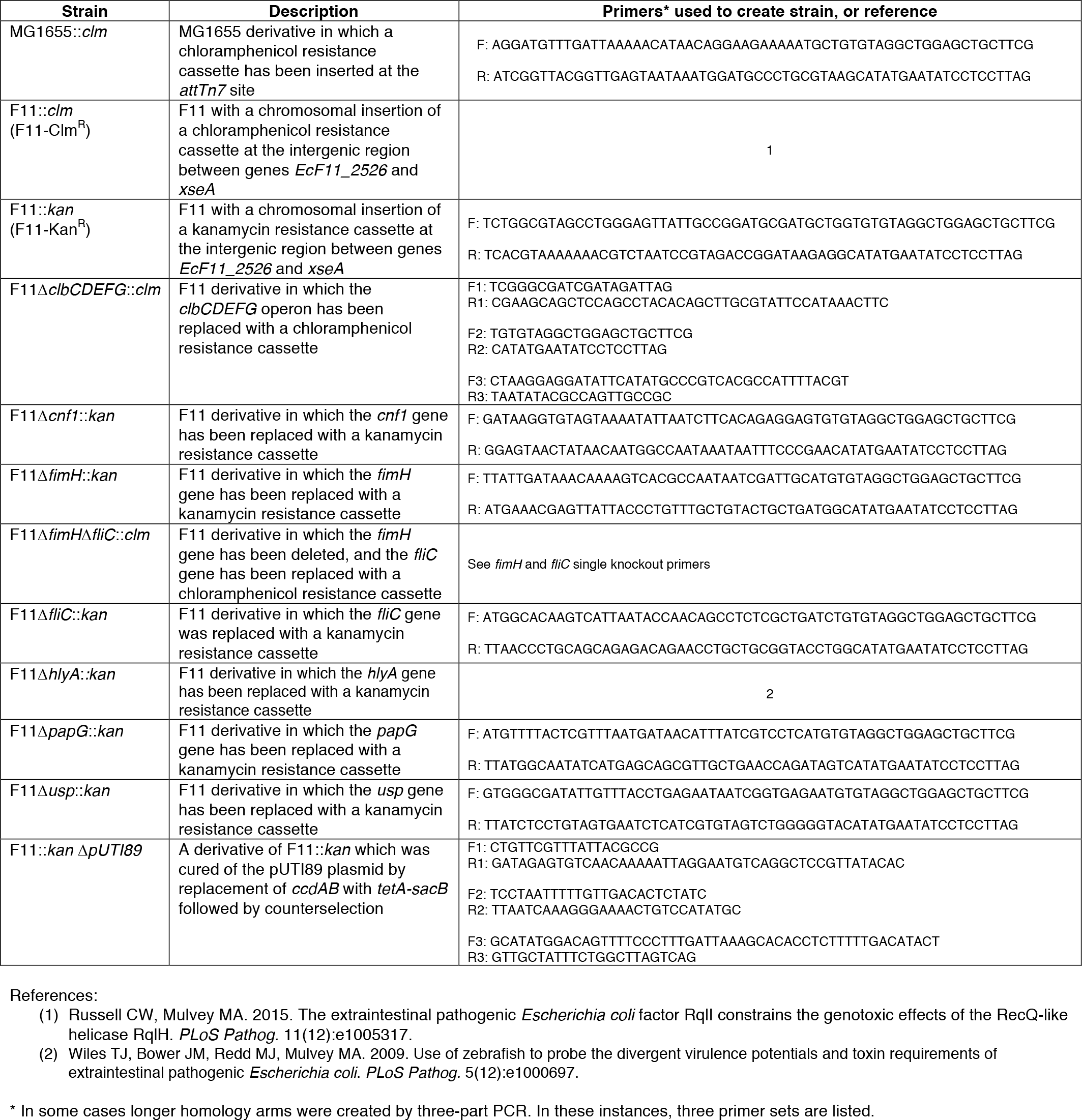
Bacterial strains used in this study.

